# New formula and conversion factor to compute tree species basic wood density from a global wood technology database

**DOI:** 10.1101/274068

**Authors:** Ghislain Vieilledent, Fabian Jörg Fischer, Jérôme Chave, Daniel Guibal, Patrick Langbour, Jean Gérard

## Abstract

**Premise of the study:** Basic wood density is an important ecological trait for woody plants. It is used to characterize species performance and fitness in community ecology, and to compute tree and forest biomass in carbon cycle studies. While wood density has been historically measured at 12% moisture, it is convenient for ecological purposes to convert this measure to basic wood density, i.e. the ratio of dry mass over green volume. Basic wood density can then be used to compute tree dry biomass from living tree volume.

**Methods:** Here, we derive a new, exact formula to compute the basic wood density *D*_*b*_ from the density at moisture content *w* denoted D_*w*_, the fibre saturation point *S*, and the volumetric shrinkage coefficient *R*. We estimated a new conversion factor using a global wood technology database where values to use this formula are available for 4022 trees collected in 64 countries (mostly tropical) and representing 872 species.

**Key results:** We show that previous conversion factors used to convert densities at 12% moisture into basic wood densities are inconsistent. Based on theory and data, we found that basic wood density could be inferred from the density at 12% moisture using the following formula: *D*_*b*_ = 0.828*D*_122_. This value of 0.828 provides basic wood density estimates 4-5% smaller than values inferred from previous conversion factors.

**Conclusions:** This new conversion factor should be used to derive basic wood densities in global wood density databases. This would prevent overestimating global forest carbon stocks and allow predicting better tree species community dynamics from wood density.

## INTRODUCTION

Wood density of woody plants is a key functional trait (Chave *et al.*, 2009; Violle *et al.*, 2007). It helps understand the functionning of forest ecosystems both in terms of carbon sequestration (Chave *et al.*, 2005; Vieilledent *et al.*, 2012) and community dynamics (Diaz *et al.*, 2016; Kunstler *et al.*, 2016; Westoby & Wright, 2006). In carbon cycle research, tree wood density is used to compute forest carbon stock and assess the role of forests in mitigating climate change (Pan *et al.*, 2011; Vieilledent *et al.*, 2016) or evaluate the impact of deforestation on climate (Achard *et al.*, 2014). In community ecology, wood density is a proxy for species performance (Lachenbruch & McCulloh, 2014), reflecting a trade-off between growth potential and mortality risk from biomechanical or hydraulic failure (Diaz *et al.*, 2016). Fast-growing, short-lived species tend to have a lower wood density while slow-growing, long-lived species tend to have a higher wood density (Chave *et al.*, 2009; Greenwood *et al.*, 2017). In wood technology, most physical and mechanical properties of wood (strength, stiffness, porosity, heat transmission, yield of pulp per unit volume, etc.) are closely related to wood density (Sallenave, 1955; Shmulsky & Jones, 2011; Thibaut *et al.*, 2001). This explains why wood density has been commonly measured in forestry institutes, where wood was principally studied for construction or paper making.

Wood density has been originally measured at ambient air moisture after air drying (Glass & Zelinka, 2010). Thereafter, wood density has been measured at fixed moisture content, such as 15% or 12%, this last value now being an international standard (Sallenave, 1955). In temperate countries, construction wood is at equilibrium with ambient air at an average moisture close to 12%. Wood density at 12% moisture is the ratio between the mass and volume of a wood sample at 12% moisture, and is expressed in g/cm^3^. In the past, this measure was also commonly reported in the British literature in pounds per cubic foot (1 g/cm^3^ = 62.427 lb/ft^3^) (Reyes *et al.*, 1992; Sallenave, 1971). In carbon cycle research and ecology, the most useful metric is the basic wood density, the ratio between oven-dry mass (at 0% moisture) and green volume (water-saturated wood volume) in g/cm^3^. This trait is sometimes referred to as wood specific gravity (abbreviated WSG). Both terms describe the same quantity but wood specific gravity is usually the ratio between the mass of a given volume of wood and the mass of the same volume of water, and is therefore unitless (Williamson & Wiemann, 2010). Here we use the term “basic wood density”. Basic wood density can be directly used to compute tree dry biomass and carbon stock from a standing tree volume estimated using an allometric equation (Brown, 1997; Chave *et al.*, 2005, 2014; Vieilledent *et al.*, 2012). For example, Chave *et al.* (2014) have estimated the following pantropical tree biomass allometric equation: *AGB* = 0.0673 × *(_ρ_D^2^H*)^0.976^ with *AGB* the tree dry aboveground biomass in kg, *D* the tree diameter at 1.30 m in cm, *H* the tree height in m and *ρ* the basic wood density in g/cm^3^. Tree dry biomass can then be converted to carbon stock using the IPCC default carbon fraction of 0.47 (McGroddy *et al.*, 2004).

Different methods have been used to convert measures of wood density at 12% moisture (*D*_12_), which are often available in forestry institute databases, into basic wood density (*D_b_*). Based on basic wood density data and air-dry wood density data (supposedly close to 12% moisture) for 379 tropical species or genera (Chudnoff, 1984), Reyes *et al.* (1992) have proposed a linear regression between *D_b_* and *D*_12_ (Eq. 1).

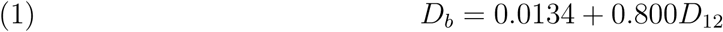

This relationship has been used to estimate the basic wood densities of 223 species in Reyes *et al.* (1992), successively reported in Brown (1997), IPCC (2006) and Zanne *et al.* (2009). Sallenave (1971) has proposed another formula to compute basic wood density from the wood density at 12% moisture (Eq. 2). In this formula, *d* is a density conversion factor per 1% change in moisture content denominated “hygroscopicity” by Sallenave (1971), S is the fibre saturation point (moisture content *S* in % at which wood volume starts decreasing in the drying process), and *v* is the variation in volume on a dry basis per 1% change in moisture content (in %/%). The values of *d, v*, and *S* vary between species and individual trees. Sallenave (1955; 1964; 1971) published values of *D*_12_, *d, v*, and *S* for 1893 trees sampled worldwide in tropical forests.

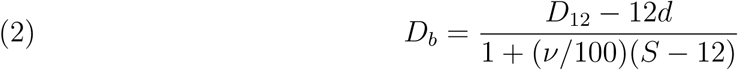

Using Sallenave’s data and formula, it is possible to compute *D*_*b,i*_ for each wood sample i and estimate the conversion factor *α*_*12*_ between wood density at 12% moisture and basic wood density from the following statistical model: *D*_*bi*_ = α_12_*D*_12,*i*_ + ε_*i*_, assuming a normal error term ε_*i*_ ~ 𝒩*ormal*(*0*, σ^*2*^). Using the wood samples of the Sallenave data-set, Chave et *al.* (2006) obtained a value of 0.872 for the conversion factor *α*_*12*_ between D_12_ and *D*_*b*_. Several studies have since used Sallenave’s method to derive conversion factors for particular sets of species (Muller-Landau, 2004) or to convert wood density at a particular moisture content *w* into basic wood density (Bastin et *al.*, 2015; Chave et *al.*, 2009; Swenson & Enquist, 2007) by extending Sallenave’s original formula assuming that *D*_*b*_ = *(D*_*w*_ *— wd)/(1* + (*v*/100)(*S* — *w*)). The resulting conversion factor was close to 0.872. Notably, Chave et *al.* (2009) used a value of 0.861 (see supplementary material of the cited reference) to convert any wood density between 10-18% moisture content into basic wood density. The estimated basic wood densities were included in the Global Wood Density Database, a large global compilation of wood density data (Chave et *al.*, 2009; Zanne et *al.*, 2009). This database combines measured (40% of the data) and inferred (60% of the data) basic wood densities. It has been extensively used to compute forest biomass and carbon stock with the aim of studying the role of forest in the global carbon cycle (Avitabile *et al.*, 2016; Baccini *et al.*, 2012, 2017; Saatchi *et al.*, 2011; Vieilledent *et al.*, 2016) or addressing questions in functional ecology (Baraloto *et al.*, 2010; Chave *et al.*, 2009; Kunstler *et al.*, 2016).

Simpson (1993) proposed a simplified formula to compute wood density at any moisture content from basic wood density. With this formula, the relationship only depends on the moisture content *w*: *D*_*w*_ = *D_b_*(1 + *w*/100)/(1 — 0.265α*D*_*b*_), with a =1 — *w*/30. Simpson’s formula can be inverted to compute *D_b_* from D_*w*_ (Eq. 3).

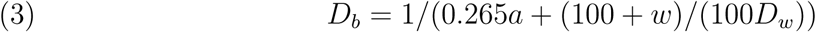

Two assumptions were made to derive this formula, (i) the fibre saturation point *S* can be approximated to 30% for all tree species, and (ii) the total volumetric shrinkage *R_T_* (in %) from *S* to 0% moisture content is proportional to the basic wood density *D*_*b*_, and can be approximated by the following relationship (Stamm, 1964): *R_T_*/100 = 0.265*D_b_*.

Because relationships proposed by Reyes *et al.* (1992), Sallenave (1971) and Simpson (1993) give significantly different estimates of the basic wood density for a same value of wood density at 12% moisture, it is important to further test their underlying theories.

In this study, we present a new and exact formula to convert wood density at any moisture content into basic wood density. The formula is derived from the definitions of the fibre saturation point and the volumetric shrinkage coefficient. We compare this new formula with formulas provided by Reyes, Sallenave and Simpson, and explain why they differ. We combine our theoretical formula with the latest version of a wood technology database compiled by Cirad (the French agricultural research and international cooperation organization) to estimate a new conversion factor between density at 12% moisture and basic wood density. We finally discuss the consequences of this new conversion factor in carbon cycle research and ecology.

## MATERIALS AND METHODS

### The Cirad wood technology database

#### A global database including 872 tree species

The Cirad wood technology database includes data from 4022 trees. Tree species names (latin binomial) were first spell-checked with the Global Names Resolver available in the **taxize** R package (Chamberlain & Szöcs, 2013) using The Encyclopedia of Life, The International Plant Names Index, and the Tropicos databases as references. Then, we searched for synonyms in the list of species names and corrected the species names when necessary using The Plant List version 1.1 (http://www.theplantlist.org) as reference. We used the **Taxonstand** R package (Cayuela *et al.*, 2017) to do so. Taxonomic families were retrieved from updated species names using The Plant List. Trees belong to 1010 taxa from 484 genera and 94 taxonomic families. Most of the taxa (872) were identified up to the species level, with varieties and subspecies combined. Out of the 872 species names, 832 were “accepted” species names and 40 were “unresolved”, according to The Plant List. The rest of the taxa (138) were identified up to the genus level. The dataset includes 834 angiosperm species and 38 gymnosperm species. The dataset includes trees from 64 countries but the major part of the trees come from 13 tropical countries (countries with more than 20 tree species), mostly in South America, Africa, and in Oceanic islands (Table 1 and Fig. 1). Sallenave was working for a tropical forestry institute now part of Cirad, the database is thus the direct continuation and extension of Sallenave’s work (1955; 1964; 1971).

**Figure 1:**
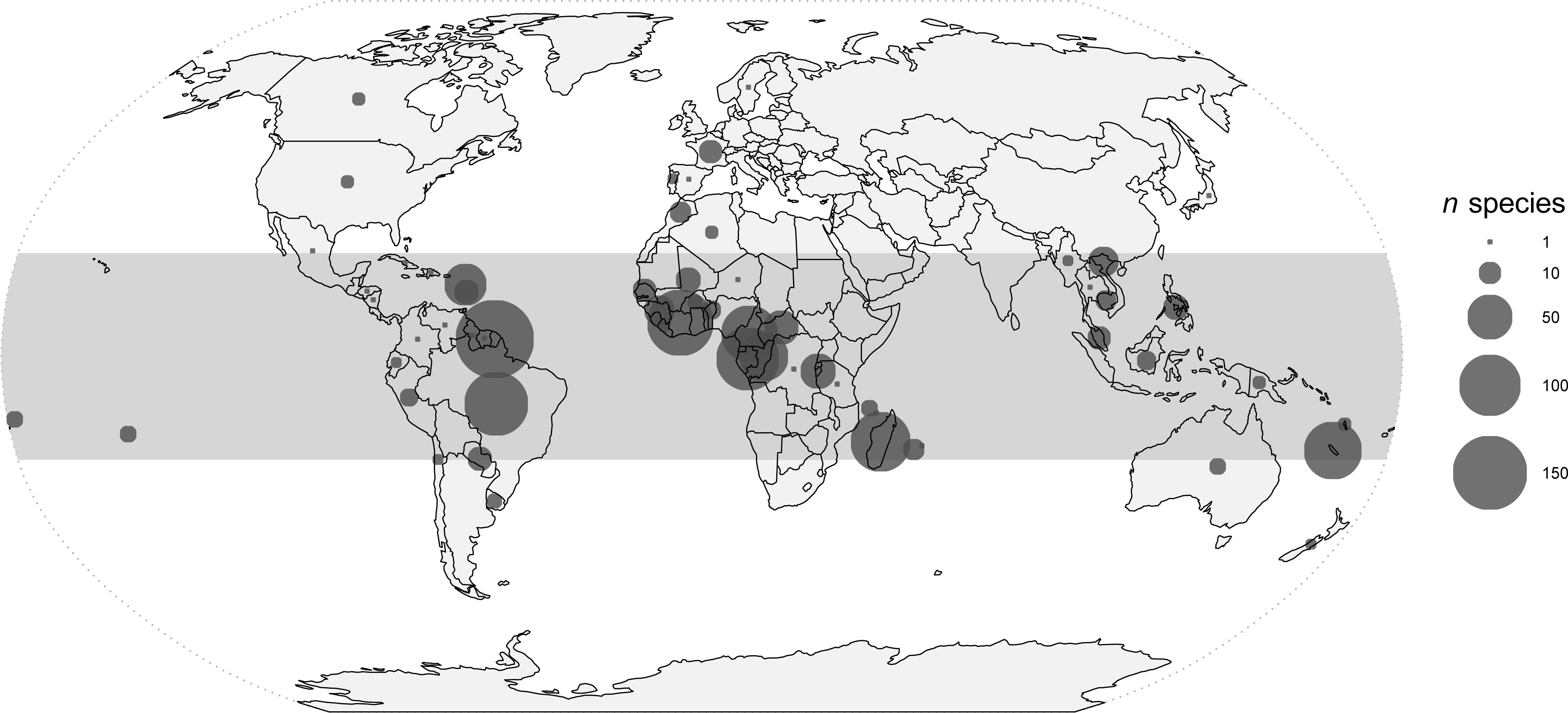
Global repartition of the data available in the Cirad wood density database. Data repartition is provided in number of species per country. Most of the species in the database (830/872) are found in the tropics (materialized by the grey band on the map).

**Table 1:** Countries with the highest number of tree species (>20) in the Cirad wood density database. The dataset includes values from 64 countries but the major part of the measurements of wood physical and mechanical properties has been done in tropical countries in South America, Africa and tropical Oceanic islands.

| Country | *n* species |
| --- | --- |
| **South America** |  |
| Brazil | 108 |
| French Guiana | 168 |
| **Africa** |  |
| Burundi | 29 |
| Cameroon | 83 |
| Central African Republic | 27 |
| Côte d'Ivoire | 117 |
| Congo–Brazzaville | 59 |
| Gabon | 105 |
| Guinea | 20 |
| **Asia** |  |
| Viet Nam | 20 |
| **Oceanic islands** |  |
| Guadeloupe | 43 |
| Madagascar | 94 |
| New Caledonia | 87 |

#### Measuring wood mass, moisture content, and volume

The volume *V*_*w*_ and mass *m*_*w*_ of a wood sample depend on its water content *w*. The moisture content of wood is a function of both relative humidity and temperature of ambient air (Glass & Zelinka, 2010; Hailwood & Horrobin, 1946). In the Cirad database, wood volume and mass measurements were done in the same laboratory following the French standard AFNOR NF B51-005 (09/1985). Wood samples are cubes of about 20 mm side (± 0.5 mm). To measure *V*_*w*_ and *m*_*w*_, wood samples were put under controled and fixed atmospheric conditions to reach a water content *w*. Wood samples were supposed to be stabilized when their variation in mass (in g) after four hours was less than 0.5%.

Wood mass *m*_*w*_ (in g) was measured with a 0.01 g precision balance. The exact moisture content *w* (in %) of a wood sample is defined as a percentage of the dry mass, *w* = 100(*m*_*w*_ — *m*_0_)/*m*_0_, with *m*_0_ being the mass of the wood sample at the anhydrous state and mw being the mass of the wood at moisture content *w*.

Wood volume *V*_*w*_ (in cm^3^) was measured with three different methods. For wood samples of irregular dimensions, we used a mercury volumenometer, or the water displacement method based on Archimede’s principle (Williamson & Wiemann, 2010). The mercury volumenometer for volume measurement was progressively abandoned from the end of years 90s due to mercury toxicity. For perfectly rectangular parallelepiped or cubic wood samples, a stereometric method was used to measure the wood cube size in the three dimensions using a digital caliper having a 0.02 mm precision. Using one of these three methods, wood volume was measured with a precision <0.003 cm^3^.

#### Measuring fibre saturation point, volumetric shrinkage coefficient, and wood density at 12% moisture

The fibre saturation point S (in %) is commonly defined as the water content above which the wood volume does not increase (Skaar, 1988). Water can exist in wood as liquid water (“free” water) or water vapor in cell lumens and cavities, and as water held chemically within cell walls (“bound” water). The fibre saturation point is the point in the wood drying process at which the only remaining water is that “bound” to the cell walls. Further drying of the wood results in the strengthening of the wood fibres, and is usually accompanied by shrinkage (Skaar, 1988).

To estimate the fibre saturation point *S*, we first measured wood volume at the saturated state *V_S_* using the water displacement method. To reach a state saturated in water, whith *w* > *S*, wood samples were autoclaved, subjected to one hour of vacuum (to accelerate water impregnation) and then soaked in water during 15 hours at 5 bar pressure. Then, wood samples were stabilized at four decreasing moisture contents *w* until reaching the anhydrous state. First, wood samples were put in a stove at 30°C temperature and 85% humidity to reach a moisture content close to 18%. Second, wood samples were put in an air-conditioned room at 20^°^C temperature and 65% humidity to reach a moisture content close to 12%. Third, they were put in a stove at 20°C temperature and 50% humidity to reach a moisture content close to 9%. Fourth, they were put in a stove at 103°C to reach the anhydrous state. Wood mass *m*_*w*_ and wood volume *V*_*w*_ were measured at each of the four stabilized stages. The exact water content w at the three stabilized states previous to the anhydrous state was computed from the mass *m_w_* and the anhydrous mass m_0_. Three volumetric shrinkage values Δ*V/V* = 100(*V_S_* — *V*_*w*_)/*V_S_* were computed between the saturated state and the three other stabilized states. The fibre saturation point *S* was defined as the intercept of the linear model *w* = *S* + *b* × Δ*V/V* (Stamm, 1964). To minimise the errors in estimating *S*, only the relationships with a coefficient of determination *r*^2^ > 98% were considered.

The volumetric shrinkage coefficient *R* (in %/%) is the variation in volume per 1% change in water content. The total volumetric shrinkage *R_T_* of the wood samples from the saturated state to the anhydrous state (in %) was computed from *V_S_* and *V*_0_: *R_T_* = 100(*V_S_*—*V*_0_)/*V_S_*. Then, the volumetric shrinkage coefficient *R* (in %/%) was estimated from R_t_ and the fibre saturation point *S*: *R* = *R_T_*/*S*. This definition of the volumetric shrinkage coefficient differs from the one used in Sallenave’s work. Sallenave used the anhydrous volume *V*_0_ as the reference volume and *v* was defined as *v* = *B*/*S* with *B* = 100(*V_S_* — *V*_0_)/*V*_0_. Because this definition corresponded to wood swelling and not to wood shrinkage, it has been changed when compiling the new Cirad wood technology database. Sallenave’s *B* values were converted to *R_T_* values with the following formula derived from the definitions of *B* and *R_T_*: *R_T_* = 100(1 — 1/(*B*/100 + 1)).

Wood density at 12% moisture (*D*_12_ in g/cm^3^) was obtained computing the ratio *m*_*w*_/*V*_*w*_ with w close to 12% moisture (when wood samples were stabilized at 20°C temperature and 65% humidity). Because the moisture content *w* was not exactly 12%, densities were initially corrected using the “hygroscopicity” term *d* defined by Sallenave and the following formula *D*_12_ = *D*_*w*_ *—* (*w* — 12)*d* (Sallenave, 1971). This correction affected only the third decimal of the wood density value, so it was progressively abandonned. Given the precision of wood mass and volume measurements (see sec.), uncertainty regarding wood densities at 12% moisture for individual samples was considered to be of about 0.01 g/cm^3^.

In the Cirad database, average values for *S, R* and *D*_12_ for each tree were historically recorded using >10 wood samples taken at various positions in the trunk. Out of the 4022 trees present in the Cirad database, 190 trees had only measurements for *D*_12_, with no values for *S* and *R*. Definitions and units of wood physical and mechanical properties used in the present study are all summarized in Appendix S1 (see the Supplementary Data with this article).

### Model relating *D_w_* and *D_b_*

Using *D*_*w*_ (the wood density at moisture content *w*), *R* (the newly defined volumetric shrinkage coefficient), and *S* (the fibre saturation point), we derived a new relationship linking the basic wood density *D*_*b*_ with *D*_*w*_. We first considered the relationship bewteen *V_S_* and *V*_w_. The volumetric shrinkage coefficient *R* (variation in volume per 1% change in water content) is defined as *R* = (100Δ*V*)/(*V*Δ*w*). Let’s consider a wood sample saturated in water (*w* = *S*) that would be dried until reaching a water content *w*. The volume of the wood sample would decrease (wood shrinkage) and *R* can be written as:

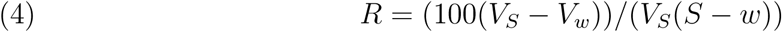

Using Eq. 4, we can express *V_S_* as a function of *V*_*w*_, *R, S* and *w*:

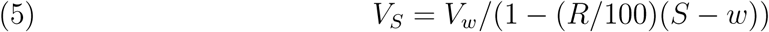

We then considered the relationship between *m*_0_ and *m_w_*. Water content *w* is defined as *w* = 100(*m*_*w*_ — *m*_0_)/*m*_0_. Using this definition, we expressed *m*_0_ as a function of *m*_*w*_ and *w*:

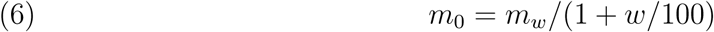

Following the definition of the basic wood density *D_b_ (D_b_* = *m*_*0*_*/V*_*S*_, *D*_*b*_), and replacing *V_S_* and *m*_0_ by their expressions in Eq. 5 and Eq. 6 respectively, we obtained *D_b_* = (*m*_*w*_/(1 + *w*/100))((1 — (*R*/100)(*S* — *w))/V*_*w*_). Given that *D*_*w*_ = *m_w_/V_w_*, we found the following relationship between *D*_*b*_ and *D*_*w*_:

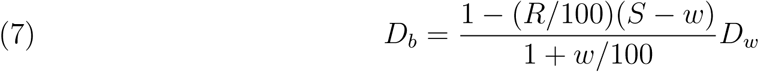

For each individual tree *i*, we used this new formula to compute the basic wood density

*D*_*b,i*_ from the values of *D*_*12,i*_ (wood density at 12% moisture), *R*_*i*_, and *S*_*i*_ reported for 3832 trees in the Cirad wood technology database (190 trees had no values for *R* or *S*). We then estimated the parameters of a statistical linear regression model linking *D*_*b,i*_ to *D*_12,*i*_ where parameter α_12_ corresponds to the conversion factor between *D*_12_ and *D*_*b*_ (Eq. 8).

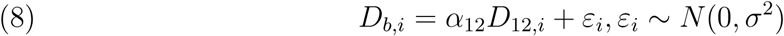

We extended this approach to compute an additional conversion factor α_15_ between *D*_15_, the wood density at 15% moisture (which was the French standard before international conventions fixed the moisture content at 12%, see Sallenave (1955)) and *D*_*b*_. We inverted Eq. 7 to compute *D*_15,*i*_ from previously computed *D_b_,_i_* values and estimated the slope of a linear regression model linking *D*_*b,i*_ to D_15,*i*_.

### Comparison with the Global Wood Density Database

The Global Wood Density Database (GWDD, http://hdl.handle.net/10255/dryad.235) provides wood densities for 8412 species from around the world (Chave et *al.*, 2009; Zanne et *al.*, 2009). The GWDD and Cirad wood density databases share common wood samples and measurements from Sallenave (1955, 1964, 1971). We quantified the amount of novel information in the Cirad wood density database. We identified and computed (i) the number of species studied by Sallenave and present in the two databases, (ii) the number of species common to the two databases but not studied by Sallenave (for which wood density values were independent), and (iii) the number of species in the Cirad database not present in the GWDD. For the species shared between databases, and with independent measurements, we compared the mean basic wood density values in the two databases. To quantify the differences between the two databases, we computed the Pearson correlation coefficient between the two values, a measure of the linear correlation (dependence), and the coefficient of variation (in %) between the two databases. The coefficient of variation is the ratio of the standard deviation of the differences between density values in the two databases divided by the mean basic wood density in the Cirad database. It is a measure of the average difference between the wood density values in the two databases. Finally, we quantified the bias (in %) in the GWDD compared to the Cirad database. This bias was defined as the mean difference between density values in the two databases divided by the mean basic wood density in the Cirad database.

## RESULTS

### Relationship between *D*_*b*_ ***and** D*_*w*_

The linear regression model linking *D*_*b*_ and *D*_12_ had a coefficient of determination *r*^2^ = 0.999 and a residual standard error of 0.015 g/cm3 (Fig. 2). We estimated a new conversion factor α_12_ = 0.828 based on the slope estimate of the linear regression. Thus, the basic wood density can be estimated from wood density at 12 % moisture from Eq. 9.

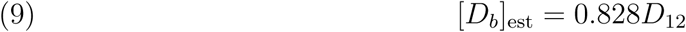

**Figure 2:**
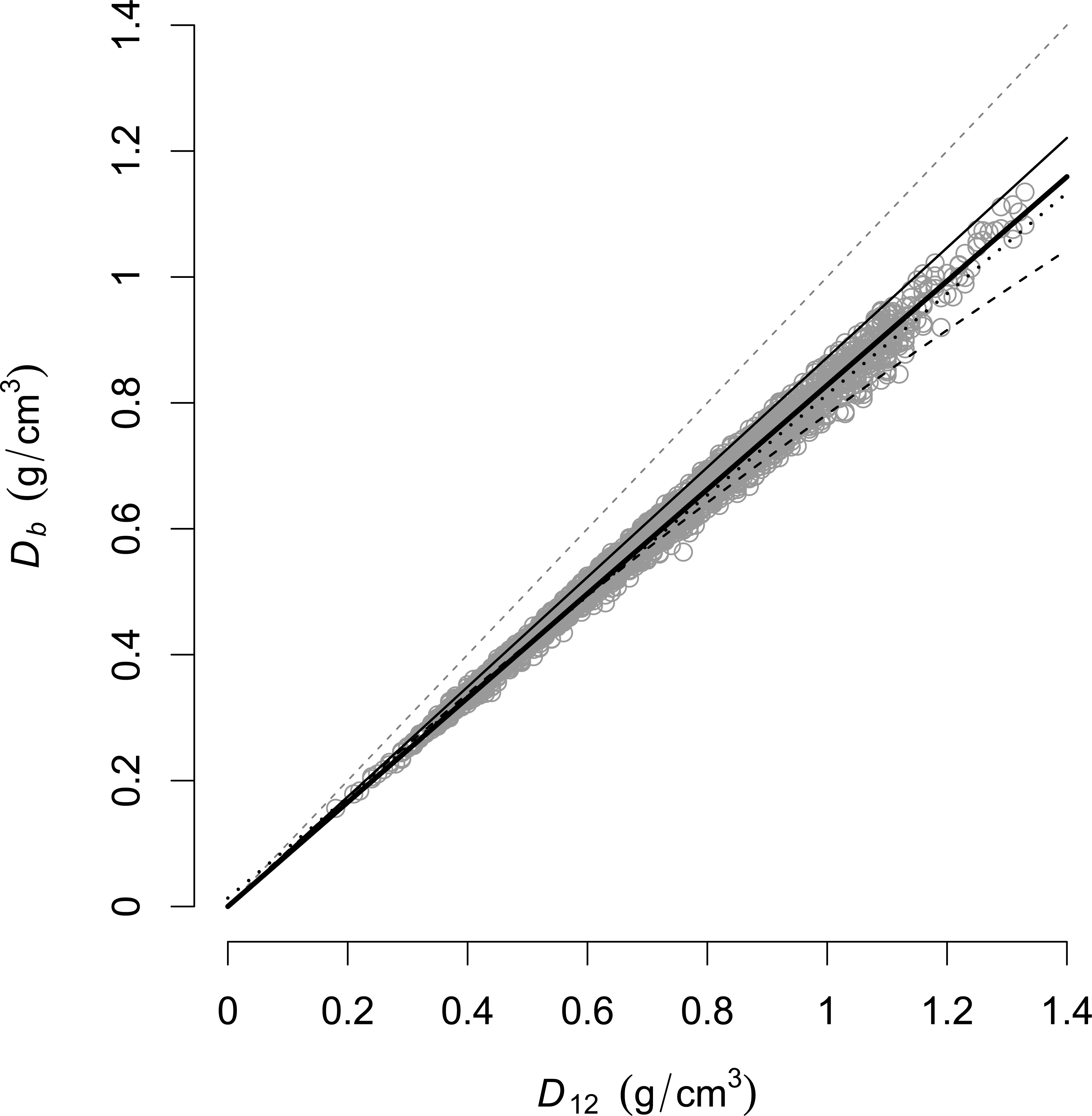
Relationship between basic wood density (*D_b_* oven dry mass/green volume, in g/cm^3^) and wood density at 12% moisture (*D*_12_). Grey dots represent the 3832 trees from the Cirad database for which *D*_12_, *R* and *S* have been measured and *D*_*b*_ computed with our new formula. The grey dashed line represents the identity line. Based on D_12_ and *D*_*b*_ values, we estimated the following relationship (plain large black line): *D*_*b*_ = 0.828 *D*_12_ (*n* = 3832, *r*^2^ = 0.999). Using Sallenave’s data and formula, Chave *et al.* (2006) estimated a significantly different conversion factor of 0.872 (plain thin black line). We also plotted Simpson’s (dashed black curve) and Reyes’ relationships (dotted black line).

With this new conversion factor, we were able to compute the basic wood density *D*_*b*_ from *D*_12_ for the 190 trees without values for *R* or *S*. At the species level, when accounting for all the trees in the data-base, *D*_*b*_ ranged from 0.191 to 1.105 g/cm^3^ (Table 2).

**Table 2:** Descriptive statistics at the species level (872 species) for the wood physical and mechanical properties in the Cirad database. See Appendix S1 for variable definitions.

| Variable | min | max | mean | median | sd | 95% quantiles |
| --- | --- | --- | --- | --- | --- | --- |
| $R$ (%/%) | 0.190 | 0.810 | 0.461 | 0.456 | 0.098 | 0.292–0.660 |
| $S$ (%) | 17 | 41 | 27.93 | 28.00 | 4.06 | 20.18–36.00 |
| $D_{12}$ (g/cm<sup>3</sup>) | 0.228 | 1.290 | 0.736 | 0.720 | 0.194 | 0.396–1.107 |
| $D_b$ (g/cm<sup>3</sup>) | 0.191 | 1.105 | 0.608 | 0.600 | 0.157 | 0.331–0.916 |

We also observed that *R, S* and *D*_12_ were not independant (Fig. 3). Thus, it is not possible to directly estimate the conversion factor from the means of *R* and *S* on the basis of the formula we derived to link basic wood density to wood density at moisture content *w* (Eq. 7). Instead, the conversion factor estimated with the linear regression model must be used.

**Figure 3:**
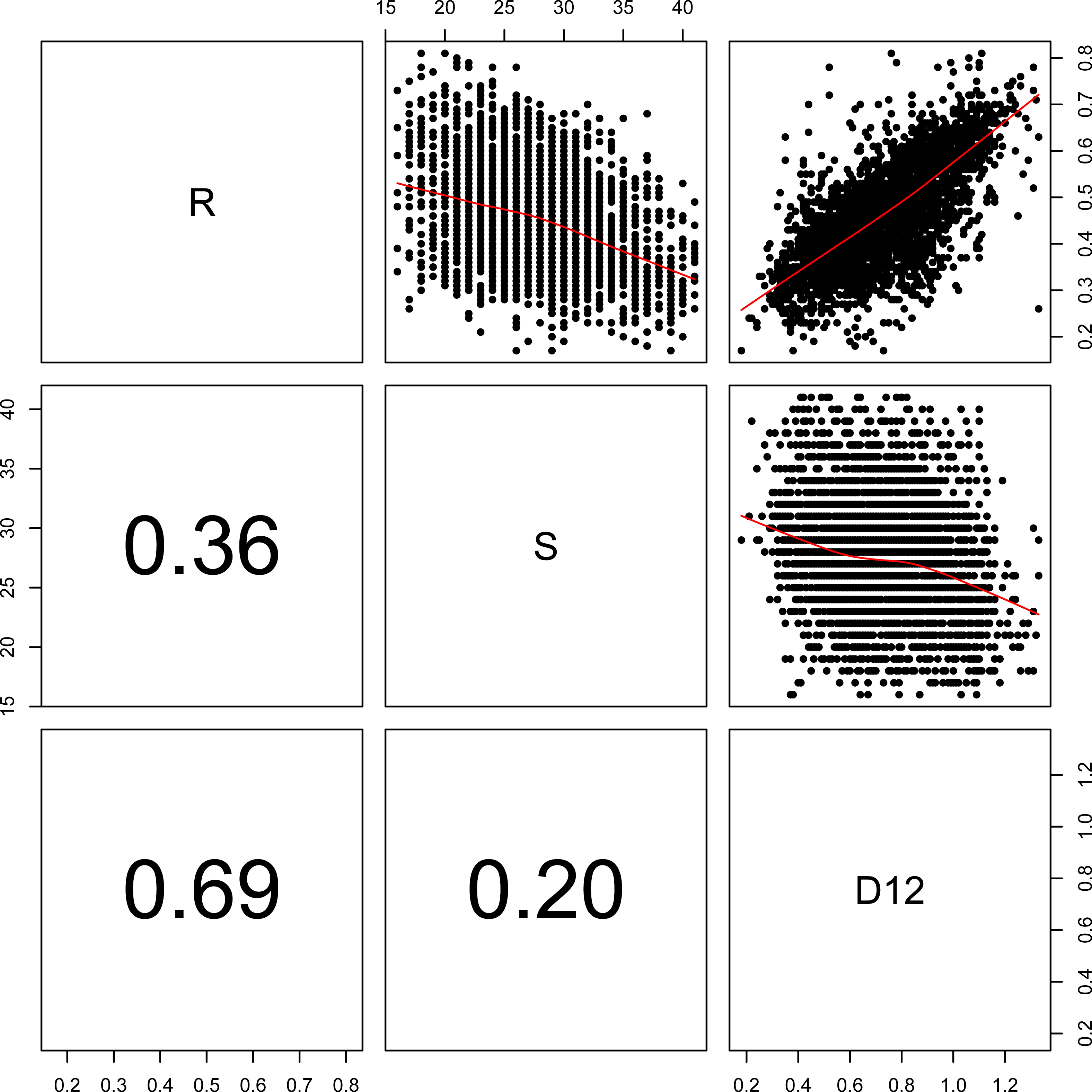
Correlation between variables describing wood properties. This figure shows the correlation between the volumetric shrinkage coefficient *R*, the fibre saturation point *S*, and the wood density at 12% moisture *D*_12_. In the lower-left panels, numbers indicate the absolute value of the Pearson’s correlation coefficient for each pair of variables. In the upper-right panels, figures show the scatter-plot for each pair of variables with a non-parametric smoother in red.

The linear regression model linking *D*_*b*_ and *D*_15_ had a coefficient of determination *r*^2^ = 0.999 and a residual standard error of 0.014 g/cm^3^. We estimated a conversion factor *α*_15_ = 0.819 between *D*_15_ and *D*_*b*_.

### Comparison with the Global Wood Density Database

Out of the 872 species in the Cirad wood density database, we identified 260 species that have been measured by Sallenave (1955, 1964, 1971) and for which one or more samples were already included in the GWDD. For these species, the Cirad database provides additional information compared to the GWDD, with values for *R, S*, and *D*_12_. We also identified 411 species common to the two databases but for which measurements of *D*_*b*_ were completely independant. For these species, the Cirad wood density database also provides *R, S*, and *D*_12_ values. Finally, we identified 201 original species in the Cirad database which were not present in the GWDD. Both *R* and *S* were highly variable among species (Table 2). In particular, *S* ranged from 17 to 41% with a mean of 27.93% and a standard deviation of 4.06%.

Using the independent measurements for the 411 common species in the two databases, we estimated a Pearson correlation coefficient of 86% and a coefficient of variation of 13.69% (Fig. 4). We also observed that, on average, *D*_*b*_ values in the GWDD were 3.05% higher compared to *D*_*b*_ values in the Cirad database.

**Figure 4:**
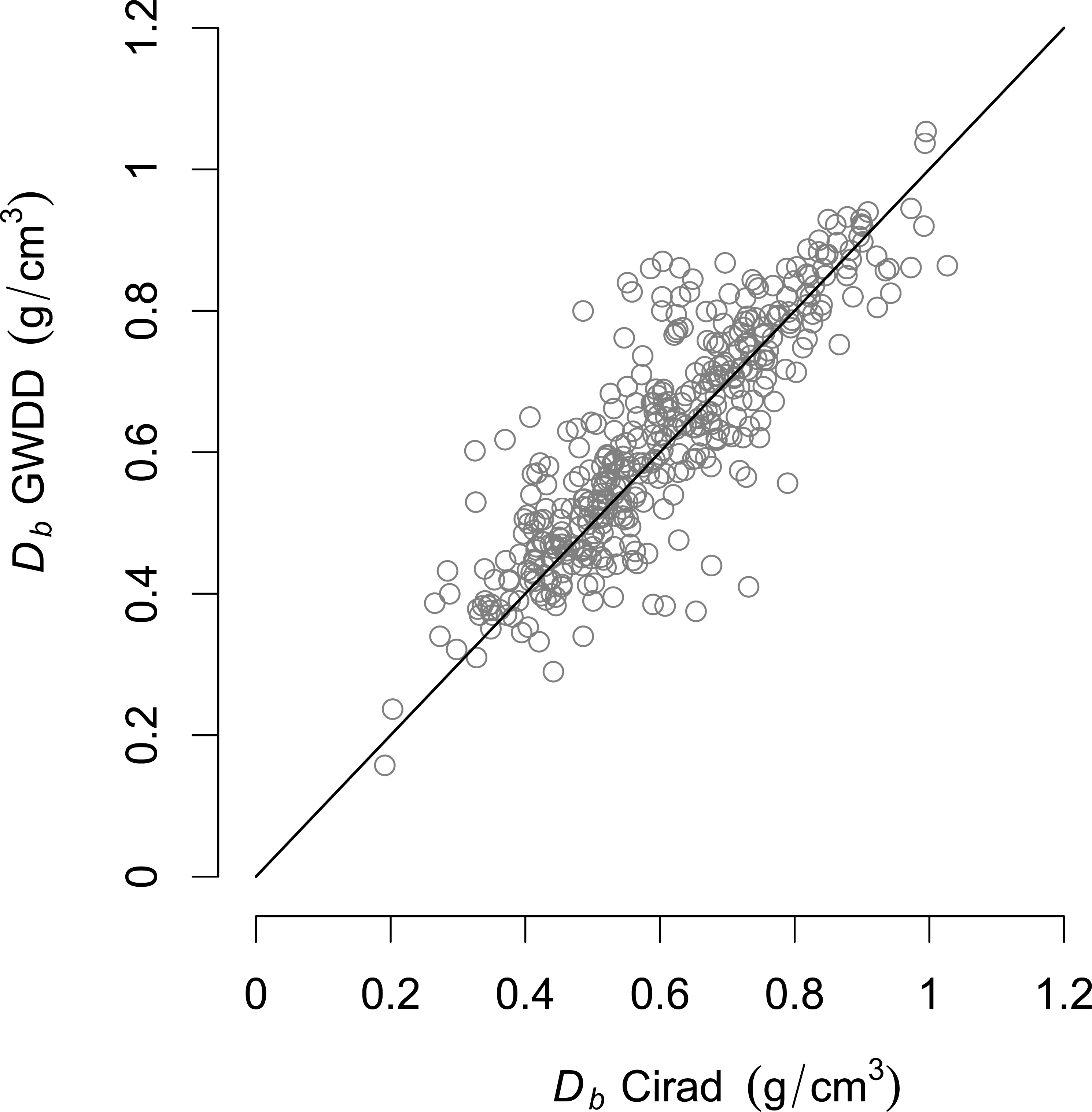
Relationship between basic wood density (*D_b_* oven dry mass/green volume, in g/cm^3^) from Cirad and GWDD databases for 411 species. The black line represents the identity line. Grey dots represent species mean basic wood densities from Cirad and GWDD databases. These 411 species are common to the two databases but wood samples and measurement protocols differ in each database. Comparing the two databases, we obtained a Pearson correlation coefficient of 86% and a coefficient of variation of 13.69%. We also observed that, on average, *D*_*b*_ values in the GWDD were higher by 3.05% compared with *D_b_* values in the Cirad database.

## DISCUSSION

### Relationship between *D_b_* and *D*_12_

We found a new value of 0.828 for the conversion factor between the wood density at 12% moisture and the basic wood density. This value is 5% lower compared to the value of 0.872 used by Chave *et al.* (2006) and based on Sallenave’s data and formula. To compare this value with the results obtained by Reyes *et al.* (1992), we derived the expectation 𝔼(*D*_*b*_/*D*_12_) from Reyes’ formula *D*_*b*_ = 0.0134 + 0.800*D*_12_. We obtained 𝔼 (*D*_*b*_/*D*_12_) = 0.0134 × 𝔼 (1/*D*_12_) + 0.800. This led to an estimate of 0.821 for the conversion factor. This value is much closer to our value of 0.828 than the value of 0.872 (Chave *et al.*, 2006).

Why was the conversion factor overestimated in Chave *et al.* (2006)? As calculations were based on the formula from Sallenave (1971), we decided to re-examine its derivation. When looking more closely at Sallenave’s own example page 11 in Sallenave (1971), a discrepancy became apparent. For the African tree species *Khaya ivorensis* (with *D*_12_ = 0.57 g/cm^3^, *d* = 0.0030, *S* = 24%, *v* = 0.46 and measured *D*_*b*_ = 0.483 g/cm^3^), Sallenave’s formula (Eq. 2) led to an estimate of 0.506 g/cm^3^ for the basic wood density. Our formula, on the other hand, gave an estimate of 0.484 g/cm^3^ which is much closer to the measured basic wood density value of 0.483 g/cm^3^. Given these findings, we suspected an error or approximation in Sallenave’s formula.

Based on the definition of the basic wood density *D*_*b*_ = *m*_0_/*V_S_* and the definition of the parameters used by Sallenave (1971), we demonstrate that Sallenave’s formula is true only if *V*_0_ = *V*_12_ (Eq. 10 and demonstration in Appendix S2). This, however, is a too strong assumption if we want to estimate an accurate conversion factor.

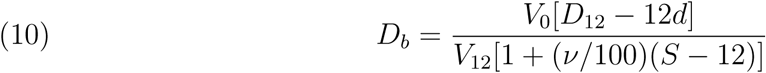

We thus recommend the use of the new formula we derived in this study (Eq. 7) to compute individual basic wood density *D*_*b*_ from *D*_12_, the wood density at 12% moisture, when *R* and *S* are available. This formula is more appropriate than Sallenave’s one. It does not only avoid making the strong assumption that *V*_0_ = *V*_12_, but also needs only two parameters to compute *D*_*b*_ compared to Sallenave’s formula which also includes a third parameter, the “hygroscopicity” d. Moreover, the new formula, unlike Sallenave’s one, implies *D*_0_ = 0 when *D*_12_ = 0, which is physically consistent. Finally, the new formula we derived in this study is more generic than Reyes’ and Sallenave’s original formula. It can be used, together with the data-set on wood properties we provide as supplementary data, to derive conversion factors between *D*_*b*_ and density *D*_*w*_ at any water content w under the fibre saturation point *S*.

We also demonstrate that our formula is more appropriate than Simpson’s one. Assumptions used to derive Simpson’s formula are not supported by our data. In the Cirad database, the fibre saturation point *S* is highly variable between species and cannot be assumed constant at 30%. We also estimated a coefficient of 0.201 for the relationship between *R_T_*/100 and *D*_*b*_, a value different from the coefficient of 0.265 suggested by Stamm (1964). We estimated a mean error (coefficient of variation of the root-mean-square-error) of 26% for *R_T_*/100 predictions, suggesting that *R_T_*/100 cannot be precisely estimated from *D*_*b*_ using a simple correlation coefficient (see also Fig. 3). As a consequence, Simpson’s formula leads to a large under-estimation of basic wood densities for *D*_*12*_ > 0.7 g/cm^3^ (Fig. 2).

If only *D*_12_ and no other measurement is available, we recommend the use of the value 0.828 for the conversion factor to compute the basic wood density *D*_*b*_. We also recommend this value of 0.828 over the value of 0.821 obtained with Reyes’ relationship. The conversion factor of 0.828 is based on a larger and more consistent database than the one used by Reyes *et al.* (1992). Database used by Reyes combined density data at the species and genera level and included air-dry densities not stabilized at 12% (Chudnoff, 1984).

### Additional value of the Cirad wood density database

Using the new formula we obtained in this study (Eq. 7), the new estimated conversion factor 0.828, and the Cirad database, we estimated the basic wood density of 4022 trees belonging to 872 species (1010 taxa), 484 genus and 94 families. Compared with the Global Wood Density Database (Zanne *et al.*, 2009), we provide basic wood density for 201 additional tree species. Most of the 872 species come from 13 oceanic tropical islands or countries.

In the Cirad wood density database, the fibre saturation point is provided for each tree. The fibre saturation point is an essential wood characteristic that can be used, in combination with the green volume, the green mass and the dry mass, to estimate the volume of water for each of the three bulk phases in a tree: (1) “free” liquid water in cell lumens and cavities, (2) water vapor in the gas-filled voids, and (3) “bound” water held chemically within cell walls (sometimes also called “solid” water, see Berry & Roderick,2005). The volume of “bound” water is an essential plant functional trait as it determines wood strength and constraints on plant architecture (Niklas, 1993), as is the volume of “free” liquid water which is the ultimate source of the biochemical activity in living plants (Berry & Roderick, 2005).

Wood characteristic values for trees in the Cirad database are the average of >10 wood samples taken at various position in the trunk. These values integrate the intra-individual variability (e.g. difference in wood density values for the same tree which can vary with the position in the trunk (Bastin *et al.*, 2015)). Providing wood characteristics for individual trees, the Cirad database can be used to compute both intra-specific and inter-specific trait variability. Intra-specific trait variability, due to genetic variability and phenotypic plasticity, participates in determining species fitness and community assemblages (Albert *et al.*, 2011; Courbaud *et al.*, 2012; Roughgarden, 1979). The Cirad database could also help quantify phylogenetic conservatism and divergences of wood densities in tree species (Flores & Coomes, 2011).

### Limits and ecological perspectives of the new conversion factor value

We found a new empirical value of 0.828 for the conversion factor. This value is obtained from a theoretical equation derived from the exact definitions of *R, S* and *D*_12_. Some uncertainty, which comes from methodological limitations associated to the measurement of these variables, is surrounding this value. In particular, the fibre saturation point S remains a theoretical concept. In practice, some “free” water is still present in wood cells when shrinkage (associated to the loss of “bound” water) starts during the drying process, and some low molecular weight organic compounds are lost during drying (Rosner *et al.*, 2009). This introduces some uncertainty in the measurement of the water content at each stage of the drying process, and thus on the estimates of *S* and *R*. Also, from the field to the laboratory, wood samples might have experienced some drying during the transport and storage, which explains why wood samples had to be re-saturated. Wood that has been re-saturated can show different shrinkage behavior (Glass & Zelinka, 2010), which introduces some uncertainty regarding the measurement of *S* and *R*. Moreover, the conversion factor could theoretically vary between species or individuals having different wood anatomies. For example, the proportion of parenchyma (representing the bulk of living cells in wood) is typically higher in angiosperms, tropical, and low wood density species than in gymnosperms, temperate, and high wood density species, respectively (Morris *et al.*, 2016). In our data-set, we found statistically significant differences between these groups of trees for the value of the conversion factor (Appendix S3). But the magnitude of the differences between groups was of the same order (≤ 0.01) as the uncertainty for the wood density value at 12% moisture *D*_12_. So we considered these differences not meaningful.

This new value of 0.828 for the conversion factor has significant implications for the study of the role of forests in the global carbon cycle. The error on the conversion factor between wood density at 12% moisture and basic wood density propagates to forest carbon stock. Combined with biomass allometric equations available in the literature (Chave *et al.*, 2005, 2014; Vieilledent *et al.*, 2012), these wood density values have been used to compute forest carbon maps globally (Avitabile *et al.*, 2016; Baccini *et al.*, 2012, 2017; Saatchi *et al.*, 2011). About 60% of the basic wood densities in the Global Wood Density Database have been estimated with an overestimated conversion factor. On the basis of 411 tree species, we showed that the GWDD overestimates wood densities by +3.05% on average. It is hard to quantify precisely the consequences of this bias on forest carbon stock estimates as it depends on relative species abundance in the forest and relative tree size distribution between species. However, if dominant species (in terms of size and abundance) have an overestimated basic wood density, due to the use of an inaccurate conversion factor (0.872 or 0.861 in Chave *et al.* (2009, 2006) against 0.828 in our study), it can potentially lead to an overestimation of 4-5% of the forest biomass and carbon stock. We are currently in the process of updating the GWDD and the present study provides a firm basis for this revision.

This study will also provide a firmer basis for future ecological research on wood density as a functional trait. Indeed, wood density is often considered as a key tree functional trait determining species performance and fitness (Baraloto *et al.*, 2010; Chave *et al.*, 2009; Diaz *et al.*, 2016; Greenwood *et al.*, 2017; Kunstler *et al.*, 2016). For example, recent global studies have demonstrated that values of wood density explained the competition outcome between pairs of tree species (Kunstler *et al.*, 2016), and that drought-induced mortality was promoted by lower wood densities (Greenwood *et al.*, 2017). Using a wood density database with unbiased values of basic wood densities would allow proper estimates of species’ differences with regards to this trait and predict better the dynamics of tree species community.

## Acknowledgments

Authors thank G. Bruce Williamson and another anonymous reviewer for their relevant remarks and constructive comments on a previous version of the manuscript. Authors warmly thank all the researchers, technicians and students who have intensively and accurately measured wood properties of thousands of trees and hundreds of species from the tropical forests at the *“Centre Technique Forestier Tropical”* and Cirad since the 1950s. They thank in particular Pierre Sallenave who made a considerable contribution to research in describing protocols and compiling data on wood properties in his three volumes (Sallenave, 1955, 1964, 1971). GV was funded by Cirad and through the European Commission ReCaREDD project at the Joint Research Center. This work has benefitted from *“Investissement d’Avenir”* grants managed by Agence Nationale de la Recherche (CEBA: ANR-10-LABX-25-01; TULIP: ANR-10-LABX-0041).

## Author Contributions

GV, FF, JC, and JG conceived the ideas and designed methodology; DG, PL, and JG collected the data; GV and FF analysed the data; GV led the writing of the manuscript. All authors contributed critically to the drafts and gave final approval for publication.

## Data Accessibility Statement

Data (including the Cirad wood density database) and R script associated to the present study have been archived on the Cirad Dataverse research data repository (http://dx.doi.org/10.18167/DVN1/KRVF0E) (Vieilledent *et al.*, 2018).

